# phuEGO: A network-based method to reconstruct active signalling pathways from phosphoproteomics datasets

**DOI:** 10.1101/2023.08.07.552249

**Authors:** Girolamo Giudice, Haoqi Chen, Evangelia Petsalaki

## Abstract

Signalling networks are critical for virtually all cell functions. Our current knowledge of cell signalling has been summarised in signalling pathway databases, which, while useful, are highly biassed towards well-studied processes, and don’t capture context specific network wiring or pathway cross-talk. Mass spectrometry-based phosphoproteomics data can provide a more unbiased view of active cell signalling processes in a given context, however, it suffers from low signal-to-noise ratio and poor reproducibility across experiments. Methods to extract active signalling signatures from such data struggle to produce unbiased and interpretable networks that can be used for hypothesis generation and designing downstream experiments.

Here we present phuEGO, which combines three-layer network propagation with ego network decomposition to provide small networks comprising active functional signalling modules. PhuEGO boosts the signal-to-noise ratio from global phosphoproteomics datasets, enriches the resulting networks for functional phosphosites and allows the improved comparison and integration across datasets. We applied phuEGO to five phosphoproteomics data sets from cell lines collected upon infection with SARS CoV2. PhuEGO was better able to identify common active functions across datasets and to point to a subnetwork enriched for known COVID-19 targets. Overall, phuEGO provides a tool to the community for the improved functional interpretation of global phosphoproteomics datasets.

## Introduction

Signalling pathways regulate the cell’s response to external perturbations and modulate some of the most important biological processes such as cell growth, differentiation, and migration^1–3^. They function through complex networks with multiple cross talks with other pathways^4–7^ and are highly context-specific; that is, signalling through the same pathway may result in completely different outputs depending on conditions, perturbations, or cell types^8–10^. Current pathway annotations don’t capture this complexity and in addition are highly biassed towards well-studied parts of the human signalling network^11,12^.

Mass spectrometry-based technologies allow us to capture in a relatively unbiased way the phosphorylation-based signalling state of a cell, through global phosphoproteomics experiments. This opens the door to data-driven extraction of condition-specific signalling networks that more accurately represent the cell’s response than existing annotated pathways.

A limitation associated with using phosphoproteomics experiments is that they are intrinsically noisy, sparse, and lack reproducibility at the peptide level^13–18^. Thus, there is a need for computational approaches that can effectively extract the active network signatures from these datasets.

One class of such methods employs network inference-based techniques, to extract a subnetwork able to explain how the phosphorylation signals propagate^19–22^. Bayesian and logic models^23–26^, ordinary differential equations^27,28^, linear and nonlinear regression^29^ and methods considering pairwise scores based on correlation^30,31^, information theory^32^ and others^33,34^, have been developed for inferring causal relationships. The HPN-DREAM network inference challenge^21^ found that the best methods typically took advantage of prior knowledge signalling pathways. This means, however, that the results from such methods often suffer from literature bias. This was evident in the inference of cell line-specific edges part of the challenge, where methods tended to perform better in cell lines that better agreed with prior knowledge networks.

Another class of algorithms, such as KSTAR^35^, KSEA^36^, IKAP^37^, KinasePA^38^, and KEA^39^, identify active kinases based on the phosphorylation levels of their substrates. However, these methods typically require a knowledge of site-specific kinase-substrate interactions, which is available only for a small number of well-studied sites. The exception is KSTAR which also accepts predicted kinase-substrate relationships. PHOTON^40^ circumvents these limitations, by integrating a set of significantly functional proteins into a protein-protein interaction (PPI) network and inferring a functionality score that is independent from the fold change of protein phosphorylation. It then uses these to derive active signalling networks from the data. However, PHOTON relies on linking ‘terminal’ nodes, i.e., the phosphorylated proteins, to a ‘source’, i.e., the receptor that was stimulated in the experiment through the ANAT method^41^. As such the results represent signalling downstream of the ‘source’ and neglect potential cross talk with other pathways or processes that might also be affected by the stimulus, but not directly linked to the ‘source’.

Recently, PPI network-based methods accounting for the global structure of the network have emerged. Distance based methods such as shortest path and network flow approaches are widely used^42–44^. Even though most of these methods are applied to transcriptomics data, they can be adapted for use on phosphorylation data. For example, PATHLINKER^45^ employs a weighted PPI network and uses a heuristic to maximise the score of the shortest paths between a set of source and target nodes. Other types of distance-based methods such as the prize-collecting Steiner tree (PCST) algorithm^46–48^ and the forest variant (PCSF)^49–51^ are also used. For example, Tuncbag et al^51^ employed the PCSF to predict multiple altered pathways in yeast from transcriptomic and proteomic data. As protein interaction networks are starting to be more systematic^52,53^, these approaches start to mitigate the literature bias issue of cell signalling studies. However, the major limitation of the distance based methods is the assumption that the shortest paths are the most informative or most likely used paths, which may not always be the case^54^.

Network propagation-based methods have been developed that boost the signal-to-noise ratio in omics datasets and predict active pathways^55^. They have been employed to predict protein functions^56,57^, prioritize candidate disease genes^58–60^, detect active modules^61–63^, and stratify patients^64,65^. TieDIE^66^ performs two propagation computations, from sources and targets, and combines the result rankings to retrieve an active subnetwork. Using this approach Drake et al^67^ extracted patient-specific network modules and potential drug targets in prostate cancer. Propagation algorithms are a perfect fit for phosphoproteomics data, which tends to be sparse, since they can fill the gap between missing values and at the same time reduce the intrinsic noise of such datasets. However, these methods do not explicitly model feedback loops, predict interaction directions, or prioritise the most likely phosphorylation regulators. Additionally, to our knowledge, they tackle the problem of signalling network reconstruction from a global perspective, but they do not consider the effect that a phosphorylated protein has on its direct functional neighbours leading to large and hard-to-interpret network signatures.

To tackle these issues, we present phuEGO, an algorithm for extracting active signalling network signatures from phosphoproteomics data. phuEGO combines a global propagation method with a local approach to extract interpretable signals from phosphoproteomics datasets and allows improved comparison and integration of datasets acquired by different groups albeit in similar conditions.

## Experimental procedures

### Datasets

We extracted the log-2 fold change of each phosphosite from the data available in http://phosfate.com68. Each phosphosite is then associated with a functional score (where available) extracted from Ochoa et al, 2019^69^. Each phosphorylated protein can be associated to multiple phosphosites and then to multiple values. To associate a single LFC and functional score to each protein we partitioned each dataset into tyrosine kinases, other kinases and phosphorylated substrates. We kept phosphorylated tyrosines and all the other kinases with a functional score and log-2 fold change (LFC) greater than the 20^th^ percentile, and we kept all the phosphorylated substrates exceeding both the 80^th^ percentile of LFC and functional score. Since multiple phosphosites from the same protein could still exceed these thresholds, we selected all phosphosites exceeding the thresholds and selected the mean LFC value and functional score, under the assumption that this would best represent the functional effect on the neighbours of the protein. This is a tuneable parameter and the user can adjust as they deem appropriate for their application.

The SARS-CoV2 datasets were extracted from the work of Higgins and colleagues^70^. In total five datasets comprising 4 different cell types at 24 hours post infection were extracted. The datasets comprise the following cell types: A549 (Higgins^70^ and Stukalov^71^), Caco-2 human lung epithelial cells (Klann^72^), Vero E6 African Green Monkey kidney cells (Bouhaddou^73^), human induced pluripotent stem cell-derived alveolar epithelial type 2 cells (iAT2, Hekman^74^).

For our analysis we selected the top 200 increased and decreased phosphosites. Where multiple phosphosites are associated with the same protein, we selected the maximum absolute LFC value.

### Pre-processing of networks and datasets

To compile the base network that phuEGO uses for its analysis we did the following: First, we retrieved the entire human protein-protein interaction network from IntAct^75^ (version: 4.2.17, last update May 2021). We also added kinase-kinase interactions and kinase-substrate interactions from PhosphoSitePlus^76^ (version 6.5.9.3, last update May 2021), OmniPath^77^ (last release May 2021) and SIGNOR 2.0^78^ (last release May 2021). Only proteins annotated in Swiss-Prot^79^ and those with at least one Gene Ontology term (GO)^80^ (last release April 2021) were retained. The resulting protein interaction network (PIN) comprises 16,407 nodes and 238,035 edges (**Supplementary Table S1**). Additionally, we modelled edge weights according to simGIC^81^ semantic similarity. The Semantic Measures Library^82^ was employed to calculate the semantic similarity among the three categories of GO (molecular function, biological process and cellular component) by adding a virtual root connecting all of them. We also generated 1,000 random networks using the configuration model available in the igraph library (method=vl). Briefly, the method^83^ implements a Markov chain Monte-Carlo algorithm to generate random networks where the node degrees are conserved. Since the edges in the random networks were reshuffled, new random interactions were created and therefore, the edge weights (i.e. simGIC semantic similarity) were updated accordingly. We applied the square (or Laplacian) normalisation to correct for the hub bias^84^. Briefly, the weight of each edge was divided by the square root of the weighted degree of the interacting nodes(1):

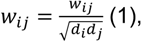

where *w_ij_* indicates the edge weight (i.e., semantic similarity) and *d_i_* and *d_j_* represent the weighted degree of node *i* and node *j* respectively.

Additionally, we also precalculated the simGIC semantic similarity of each node in the PIN against all the other nodes to associate the mean and the standard deviation to each node in the PIN. These values are used in next steps of the method to filter the ego networks by calculating the z-score (see paragraph on ego decomposition below).

### Network propagation by random-walk-with-restart

PhuEGO accepts as input a dataset of phosphorylated UniProtKB entries and the corresponding log-2-fold change (LFC). PhuEGO first assesses the prior seed set of nodes, meaning the nodes in the PIN from where random walkers should start. To do so, the input dataset is initially divided into positive and negative LFC. These two partitions are subsequently divided into: (i) the tyrosine kinases, (ii) the rest of kinases and (iii) the non-kinase phosphorylated proteins. To assess which proteins will be assigned to each partition we retrieved all the human kinases (Clan CL0016) from the Pfam^85^ database (Pfam ver 34.0 released in March 2021). Since we wanted to distinguish the tyrosine kinases from the rest of kinases, we retrieved all the human tyrosine kinases associated with the Pfam domain (PF07714) from UniprotKB^79^. In total, 531 kinases are present in our PIN, of which 127 are tyrosine kinases (**Supplementary Table S1**).

Each of the three partitions corresponds to three different restart probability vectors, whose dimension is equal to the number of nodes in the PIN and the restart probability values are equal to the LFC of the phosphorylated proteins, scaled between 0 and 1. Therefore, we start three distinct RWR runs, one for each partition, and each one involving different sets of prior nodes. As a result, we obtain three probability vectors, one for each partition, representing the most probable nodes from the perspective of the seed nodes. The idea behind this procedure is to mimic signal propagation in a global manner having as central input nodes the phosphorylated proteins and integrating the signal from these with that from the kinases, as the drivers of cell signalling. To assess the significance of the resulting RWR values we repeated the same procedure using the same seed nodes but against 1,000 random networks, generated as described above. This allows us to estimate the empirical P-value for each node of the network as the percent of its random scores that exceed the real score. At the end of this process only the nodes with a P-value<0.05 are maintained independently of which partition they have been assessed with. Note that this process is repeated two times, one for the upregulated phosphoproteins and one for the downregulated ones; as a consequence, two subnetworks are extracted associated with increased and decreased phosphorylation levels respectively.

### Generation of functional ego networks

We extract ego networks as a subgraph centred on a seed/phosphorylated node and comprising all the overrepresented nodes in a 2-steps distance from the ego. Since ego networks are still highly interconnected, in theory they could have the same dimension as the signalling network from the initial random-walk-with-restart process. To select only those ego neighbours that are most functionally similar to the ego, we computed the z-score associated with each ego neighbour using the precomputed mean and the standard deviation of the simGIC score (see Preprocessing of networks and datasets paragraph). Only those with a z-score>1.64 are selected (2)

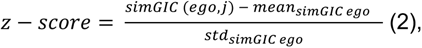

where the *simGIC(ego,j)* is the semantic similarity between the ego and node *j*, and the mean and std are the mean and the standard deviation of all the semantic similarity between the ego and all the other nodes of the PIN. The nodes with z-score>1.64 represent the functional ego network since they are also the most similar in terms of semantic similarity to the GO terms in which the ego is involved in. Let Γ(ego) represent the 1^st^ order neighbours of the ego node and Γ_Γ(ego)_ the 2^nd^ order neighbours of the ego network. The edge weights are updated according to (3):

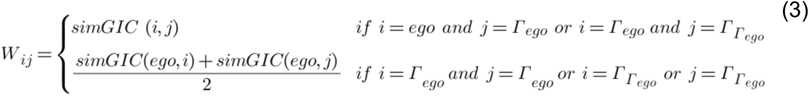

The ego networks obtained are normalised to correct for hubs using the Laplacian normalisation as in (1) (**Supplementary Figure 1A&B**).

### Ego decomposition

To understand which nodes are more closely related to the ego and, hence, involved in a similar process/pathway, we decomposed each ego network with greater than 5 neighbours into two vectors, one representing the topological distance from the ego, and one the functional distance from the ego.

To calculate the topological proximity, each node of the ego network is the source of a second run of RWR with a damping factor equal to 0.85. The restart probability vector is filled with 0 except in the node under consideration which is equal to 1. To calculate the distance between the ego node and all the other nodes of the ego network, the following formula is used (4):

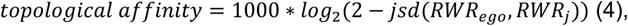

where *jsd* refers to the Jensen-Shannon distance, representing the similarity between two probability distributions. The RWR_j_ refers to the RWR probability vector when one of the nodes of the ego network is selected as seed, the RWR_ego_ refers to the RWR probability vector when the ego is the seed node. Nodes with values close to 1 are considered topologically similar to the ego.

The functional vector is defined as the logarithm of the semantic similarity between the ego and any other nodes in the network (5).

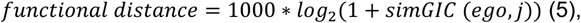

where *simGIC* represents the semantic similarity measure between the ego and the node *j*.

To identify the most similar nodes to the ego we employed the Kernel Density Estimation (KDE) using the Gaussian kernel, where each node is represented as a point in a 2D plane where the x-axis represents the topological affinity to the ego and the y-axis represents the functional similarity to the ego (**Supplementary Figure 1C**). The bandwidth for the KDE is estimated using the Silvemann formula^86^. KDE estimates the joint probability density function of the topological and semantic similarity vectors obtained at the previous step. We then calculated the joint cumulative distribution function and select only those nodes according to the following formula:

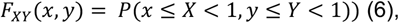

where x and y are user defined parameters. For this paper, we set these parameters to 0.85 or 0.9 depending on the application (see respective sections).

### Defining the supernode network through merging the ego networks

Each ego node and the neighbours exceeding the user’s selected probability threshold, constitutes a supernode, i.e., a small cluster of proteins that are topologically and functionally related to the ego and therefore potentially affected by its phosphorylation. We then calculated the relationships between all the supernodes to generate the supernode network. To do so for each combination of two ego nodes, we extracted the subnetwork originated by the union of the nodes included in the supernode pair and normalised it according to (1). The two egos, if connected, represent the sources of a third RWR run with a damping factor equal to 0.85. We calculate the weight between supernodes using (7)

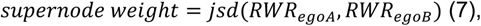

where the RWR_egoA_ refers to the RWR probability vector when the ego_A_ is selected as seed, the RWR_egoB_ refers to the RWR probability vector when the ego_B_ is selected as seed node. Edge values close to 0 indicate a strong relationship between supernodes, meaning that they potentially share many neighbours. Note that the link between two supernodes is not necessarily associated with a physical interaction. We then applied the Leiden^87^ algorithm to the supernodes network to extract functional modules. Note that the Leiden algorithm is only applied to all the connected components bigger or equal to 4 supernodes. The connected components containing less than 3 supernodes are considered as functional modules (**Supplementary Figure S2**). Isolated supernodes are removed.

### Evaluation through enrichment analysis and overlapping coefficients

Enrichment analysis is a standard approach employed to determine if known biological functions or processes are over-represented (enriched) in a set of genes/proteins of interest. The enrichment analysis is based on Fisher’s exact test which assumes that the data is hypergeometrically distributed. We used the nodes in a module as a foreground for the enrichment analysis while the human PIN is used as the background. Additionally, the P-values obtained are Bonferroni corrected. Enrichment analysis can be performed against several databases such as GO, KEGG^88^, Reactome^89^, Bioplanet ^90^, DisGeNET^91^.

To assess the similarity between modules and the reference pathways we employed the overlap coefficient or Szymkiewicz–Simpson coefficient (8).

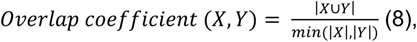

where *X* and *Y* represents the two sets of proteins under consideration. We also measured the pairwise overlap distance^92^ between the following KEGG reference pathways: Cell cycle, EGF, TCR, MAPK, VEGF, TGF, Insulin, NGF, by employing this formula:

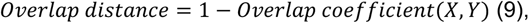

where X and Y represent the set of proteins involved in the respective reference pathways. Therefore, the distance between a pathway and itself is equal to 0. Since phuEGO extracts on average 4 modules from each dataset, modules comprising less than 10 proteins are discarded to avoid increasing the overlapping coefficient artificially with very small modules against very large ones. Additionally, we selected the modules with the best overlapping coefficient regardless of the pathway they could be annotated with.

The performance of phuEGO was compared to the enrichments resulting from a) the seeds b) the network resulting from the initial RWR step and c) the Prize Collecting Steiner Forest algorithm (PCSF) from the omicsintegrator2 package^93^. In brief, PCSF works by identifying an optimal forest in a network by maximizing the collected prizes and minimizing the edge costs. We performed a grid search for each dataset to fine tune the parameters to select the best network from Omicsintegrator2. We selected the following parameter ranges *ω* = [0.25, 0.5, 0.75, 1], β= [0.25, 0.5, 0.75, 1, 1.5, 2], *γ*=[3, 3.5, 4, 4.5] and selected the network with the best objective function. In particular, *ω* regulates the number of selected outgoing edges from the root, β is a scaling factor of prizes, and *γ* controls the edge penalty on hubs.

### Comparisons of SARS-CoV2 networks and seed nodes

To compare the networks generated by phuEGO from Higgins et al. and the seeds, we assigned to each node in the corresponding network the average RWR values from each of the three partitions or 0 if not present. We performed the same procedure to compare the seeds nodes alone with the exception that we employed the LFC values. Then we used the *hclust*package (https://www.rdocumentation.org/packages/stats/versions/3.6.2/topics/hclust) to perform the hierarchical clustering and *dendsort* (https://cran.rstudio.com/web/packages/dendsort/index.html) to optimize the ordering of leaves in the dendrogram.

### Enrichment of known targets in SARS-CoV2 datasets

Known targets for COVID-19 were extracted from Open Targets (February 2023) using the query ‘MONDO_0100096’. In total 390 drug targets were extracted, of which 365 were present in the network. Only the SARS-CoV2 networks with a damping factor equal to 0.85 and a KDE threshold>=0.85 were selected (**Supplementary Table S2**). To generate the A549 SARS-CoV2 network we selected the nodes in common between the Higgins^70^ and Stukalov^71^ network. To assess the overlap between the network nodes and the known targets we used Fisher’s exact test considering as background the entire PIN used in the analysis.

### Data visualisation

Plots were generated in Python v3.10 using the *seabron* and *matplotlib* libraries. Cytoscape (v3.9.1) was used for visualising networks. Enrichment maps were generated in R with the *clusterplofiler* and *enrichplot* packages. Hierarchical clustering of the SARS-cov2 datasets was done in R with *pheatmap*.

### Experimental Design and Statistical Rationale

The datasets included in this study were selected as they formed a unified collection by Ochoa and colleagues and included a diverse array of stimulations including well-studied and less commonly studied pathways. They were therefore available both individually from the original studies and as a reanalysed uniform set, and we could therefore compare the performance on both sets. There was also a functional score annotation available for most peptides included, allowing us to evaluate the ability of phuEGO to improve the signal-to-noise ratio. The p-values of the nodes included in the final ‘global’ networks were calculated as described above and the choice of 1000 random networks for this calculation was made to allow for a resolution of three decimal points in the calculations, while keeping the computational time still reasonable. Network and hub normalisations and statistical tests were done according to common practice as described in the respective sections above.

## Results

### A method to extract signalling modules from phosphoproteomics data

We developed phuEGO, an algorithm to reconstruct active signalling pathways from phosphoproteomics data (**Figure 1**). PhuEGO comprises two steps: a) an initial filtering of a global protein interaction network (PIN) compiled from the literature (IntAct^75^, SIGNOR^78^, PhosphositePlus^76^ and OmniPath^77^; see **Experimental Procedures**) to coarsely identify networks associated with increased and decreased phosphorylation using random-walk-with-restart and b) a step to extract the local effect of each differentially abundant phosphosite on its neighbourhood, from these larger networks.

**Figure 1.**
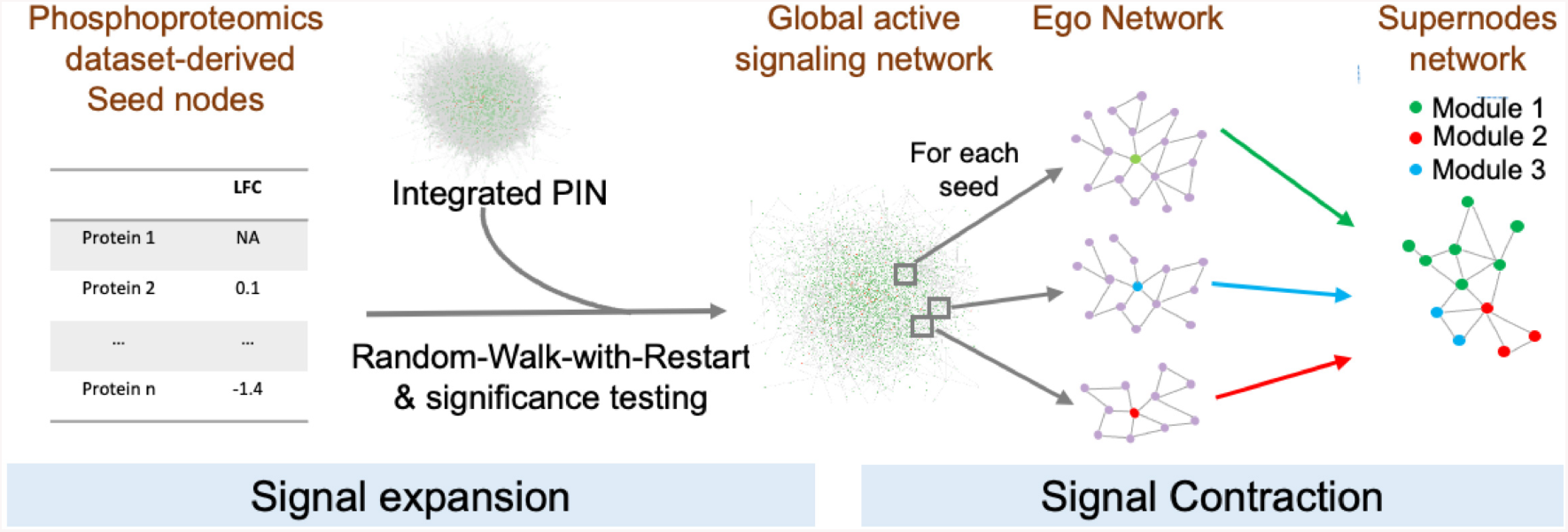
Overview of phuEGO’s methodology. PhuEGO first starts by performing a three-layer random-walk-with restart on an integrated protein interaction network. It then re-maps the seed nodes as ‘egos’ and identifies a local network or module that comprises nodes that are most topologically and functionally similar to the ego. Finally, by combining these modules phuEGO generates a network of supernodes that include overlapping functional modules.

Specifically, phuEGO first generates global networks as a result of random-walk-with-restart performed three times - from (i) tyrosine kinases, (ii) other kinases and (iii) substrates identified in the phosphoproteomics datasets (**Experimental Procedures**). This reflects our knowledge that kinases are the main drivers of phosphorylation-based signalling responses, with tyrosine kinases typically acting upstream of the global signalling response^94,95^. Upregulated and downregulated phosphosites are treated separately to uncover two networks associated with each class of phosphosites: an upregulated ‘active’ network and a downregulated one. This parameter is tuneable by the user to provide input that takes into consideration, e.g., phosphosites known to inactivate proteins.

These coarse networks comprise on average ~2,500 nodes (**Figure 1**; **Supplementary Figure S3**). To improve the interpretability of the phosphoproteomics datasets and extract more specific up/down-phosphorylated signalling modules phuEGO uses ego network decomposition to capture the functional and topological effect of the phosphosites identified in the datasets locally. Ego networks represent small subnetworks comprising all the nodes that are two steps away from the ego, which phuEGO further reduces by removing nodes that are not functionally similar to the ego (**Experimental Procedures**). By combining ego network embedding with kernel density estimation (KDE) phuEGO selects the nodes that are most similar to the ego thus generating supernodes, which are small networks comprising the ego and the functionally related neighbours. Then phuEGO generates the supernodes network where the edges weights represent the relationships between supernodes. The Leiden algorithm^87^ is employed to partition the supernode network. This procedure generates modules (~3-4 on the datasets tested in this work; **Supplementary Figure S4A**), comprising the ego and neighbours’ nodes (~25-50 nodes on average; **Supplementary Figure S4B**) that are more functional and topologically similar to the ego and, therefore, are more likely to be relevant to the signal represented by the ego. Thus, given a phosphoproteomics dataset, phuEGO extracts interpretable signalling subnetworks, associated with increased and decreased phosphorylation.

### phuEGO boosts the signal-to-noise ratio

As a first step in validating whether our method is indeed able to boost active signals from phosphoproteomics datasets we evaluated in 46 datasets (**Supplementary Table S3**), whether the phuEGO-extracted active networks were more enriched in the prior knowledge pathways that are expected based on the stimuli^88^, than the raw upregulated phosphoproteins in the datasets. We also compared the enrichment ranking to using the RWR alone (Signal expansion stage; **Figure 1**) as this would be equivalent to other methods that use network propagation to boost functional signal from omics datasets^40,66^. To our knowledge, there are no other methods that can serve a similar function as phuEGO, with the exception of PHOTON^40^, which we were unable to run as it appears to be no longer maintained. The Prize Collecting Steiner Forest algorithm (PCSF) is not a network propagation-based approach, but has been used successfully previously to identify network signatures from phosphoproteomics datasets^96^, and we therefore included it in our performance comparison.

We considered pathways as ‘more enriched’ when they ranked at a higher percentile of the total pathways found (P value<0.05, Bonferroni corrected; **Experimental Procedures; Supplementary Data S1**) in the phuEGO networks compared to the raw set of differentially abundant phosphosites (seeds). Overall, phuEGO boosts the ranking of the expected pathway for all datasets (**Figure 2A**). Where the signal is already well-defined in the seeds, it maintains the high ranking and doesn’t introduce further noise to dilute it through the diffusion process. Impressively, it is able to identify and rank highly the correct pathways even in datasets where the signal was initially very weak (e.g., Olsen et al, 2010 150 and 180 minutes) or not present at all among the seed nodes (e.g. Olsen et al, 2010 450 minutes; **Figure 2A; Supplementary Table S3**).

**Figure 2.**
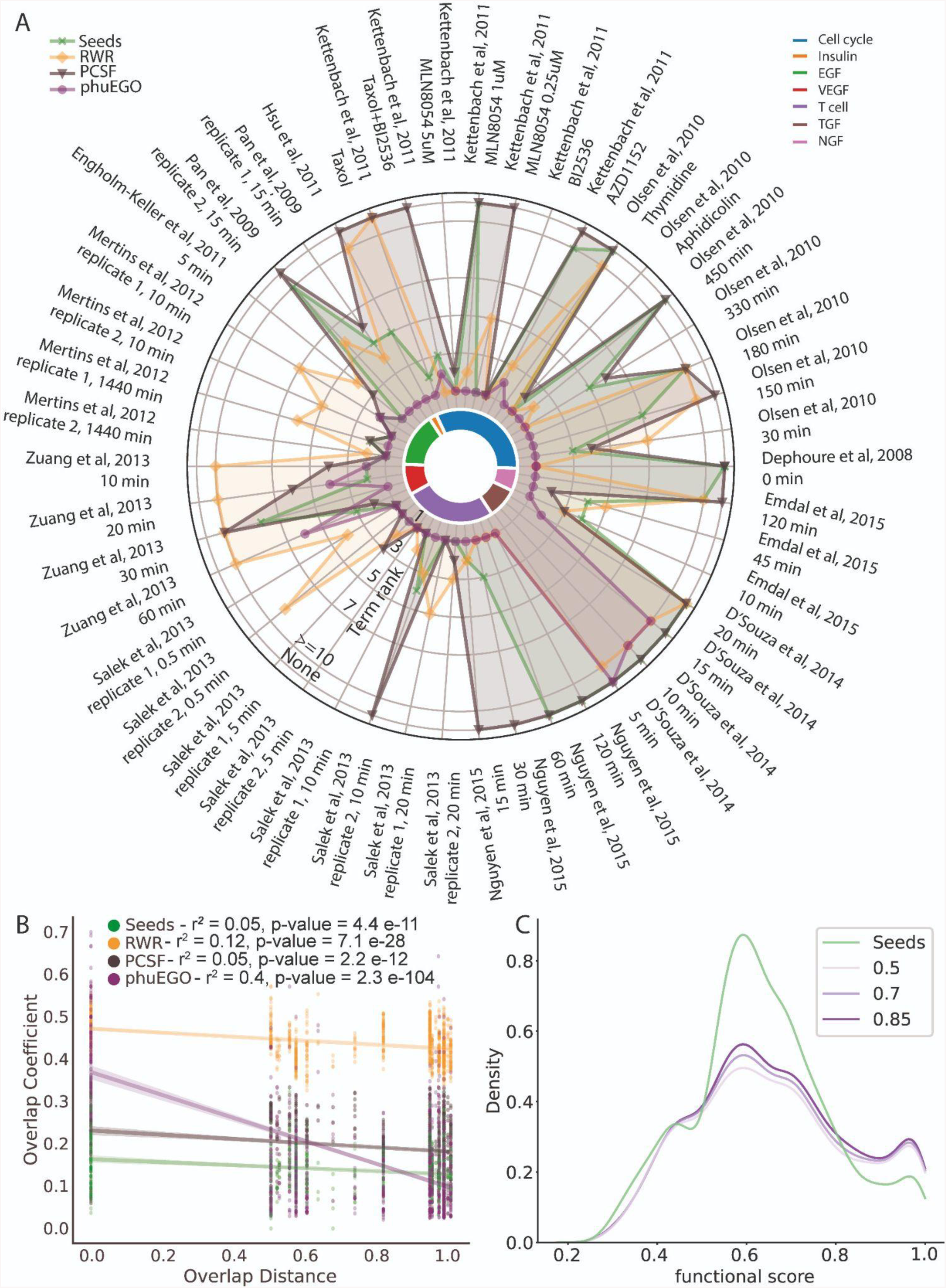
Evaluation of phuEGO. **A.** Comparison of seeds, PCSF, RWR and phuEGO with respect to their ability to rank highly the expected dominant signal, as defined by pathway enrichment analysis. The centre of the circle indicates the relevant pathway ranked first and the perimeter indicates a failure to identify the pathway at any rank. **B.** Association of the overlap coefficient of the phuEGO active signature with the overlap distance of the expected pathways across all pairwise pathway comparisons in our benchmark test. **C.** Phosphosites/nodes retained by phuEGO tend to have a higher functional score indicating an improvement in the signal-to-noise ratio of the active signatures.

When comparing to the alternative approaches (RWR and PCSF) phuEGO generally performs better, ranking the relevant enrichment term higher or similar in all but one dataset from D’Souza et al 2014 (in two out of the three time points), where even for the seeds and PCSF that performed better the ranking was very low (**Figure 2A**).

One of the main aims of our algorithm is to decrease the intrinsic noise of the phosphorylation datasets and improve their ability to identify the active signalling responses reproducibly. We thus evaluated whether phosphoproteomics experiments treated with the same conditions were more similar to each other before or after phuEGO was applied.

We first computed the overlapping coefficient between the seed nodes and the respective target pathway as the baseline. Each of the clusters that phuEGO identifies represents a unit of signalling, similar to a pathway. We don’t expect all cell lines/types to have the exact same global response to the same stimulus, but we do expect at least one of these signalling modules to be similar. We thus extracted the corresponding pathways with the highest overlapping coefficient between datasets. The similarity of these pathways should be roughly analogous to the similarity of the prior knowledge pathways that we expect to be activated with the given stimuli (**Experimental Procedures**). We found that before phuEGO there is no relationship between the overlap of the phosphoproteins and the similarity of the prior knowledge pathways that we expect to be activated (r^2^=0.05, p-value=4.4e-11). This is also true for the RWR (r^2^=0.12, p-value=7.1e-28) and PCSF (r^2^=0.05, p-value=2.2e-12) approaches (**Figure 2B**). Conversely the pathways identified by phuEGO have an overlapping coefficient that is correlated to that of the respective prior knowledge pathways (**Figure 2B**; r^2^=0.44 pvalue=8.9e-120). This is true both using the full collection of datasets as reprocessed by Ochoa et al^69^ and when using the data from the original publications (**Supplementary Figure S5**).

To assess whether phuEGO indeed is able to reduce the inherent noise of phosphoproteomics datasets we evaluated whether phosphosites that survived the process and remained as part of an integrated active signal, i.e., supernodes, had a higher functional score compared to those that remained isolated and were therefore filtered out. The functional score was extracted from Ochoa et al, 2019^69^ and ranges from 0.0 to 1.0 with higher values representing increased likelihood that the phosphosite will have a regulatory function on the protein that carries it. Across the 46 datasets (**Supplementary Table S3**) we found that phuEGO supernodes that remained as part of the active signalling signature were indeed significantly more functional than those that were filtered out (**Figure 2C**; Mann-Whitney-U pvalue (damping=0.5) =4.8e-9, Mann-Whitney-U pvalue (damping=0.7) =5.5e-10, Mann-Whitney-U pvalue (damping=0.85) =6.3e-12). Therefore, phuEGO can filter out phosphosites that are less likely to be functional and thus represent noise in the dataset.

Together these analyses demonstrate how phuEGO is able to boost the active signal, while reducing the noise in global phosphoproteomics datasets.

### phuEGO can distil the active signalling networks from diverse phosphoproteomics studies of SARS-CoV-2 infection

As a case study, we compared 5 phosphoproteomics datasets compiled from the literature by Higgins and colleagues^70^ at 24 hours post infection since this time point was common to all the datasets. The datasets are targeting different cell types: A549 (Higgins^70^ and Stukalov^71^), Caco-2 human lung epithelial cells (Klann^72^), Vero E6 African Green Monkey kidney cells (Bouhaddou^73^), human induced pluripotent stem cell-derived alveolar epithelial type 2 cells (iAT2, Hekman^74^). We found that the agreement of increased and decreased phosphorylation abundance LFC (**Figure 3A, Supplementary Figure S6A**) was very low between datasets, as measured by Pearson correlation, even when comparing experiments done on the same cell type. When phuEGO is applied (**Supplementary Data S2**), the correlation increases and is even higher when comparing the same cell types (**Figure 3B, Supplementary Figure S6B**).

**Figure 3.**
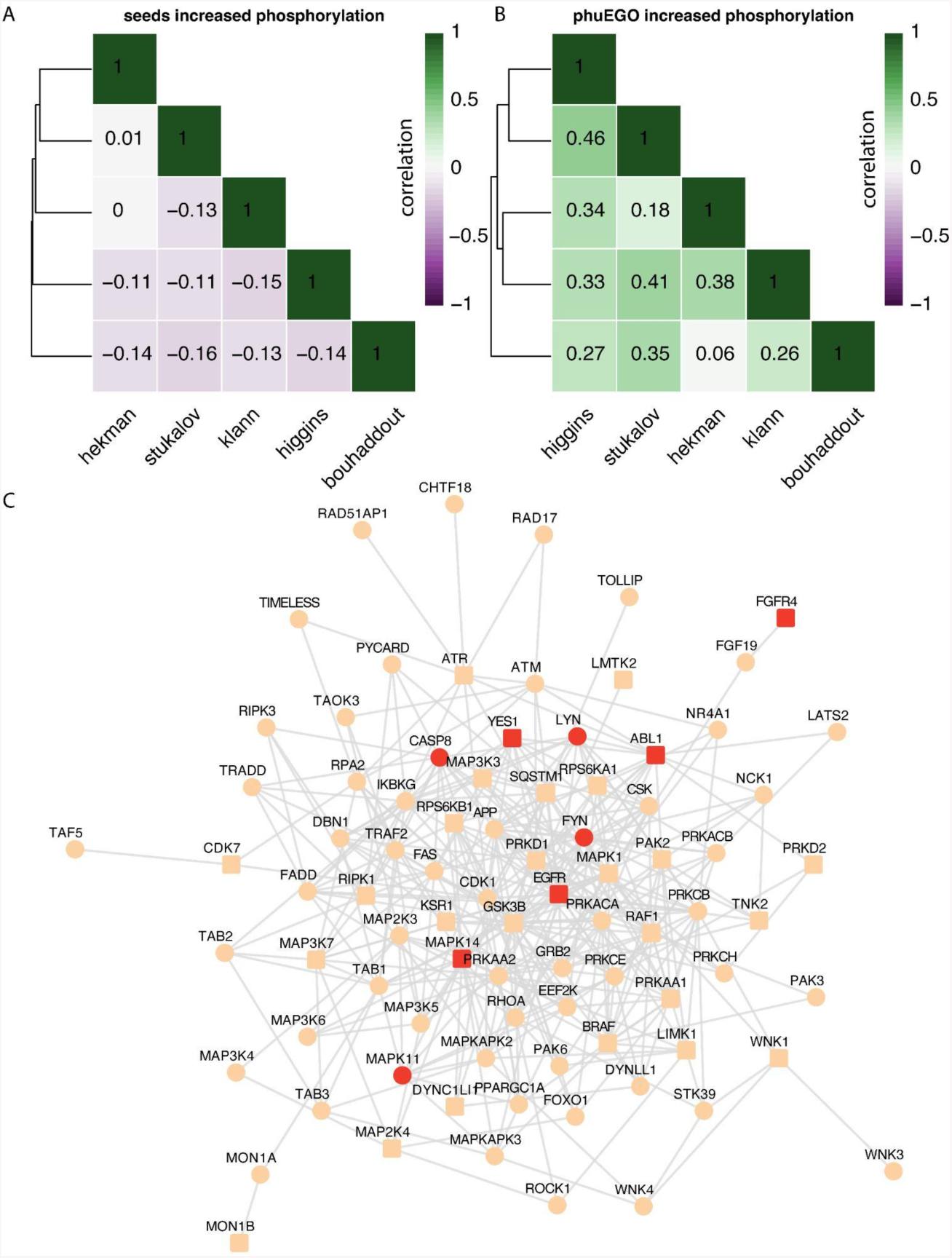
PhuEGO extracts active signatures of SARS-CoV-2. **A.** Public phosphoproteomics datasets of SARS-CoV2 infection correlate poorly. **B.** The correlation of public phosphoproteomics datasets upon SARS-CoV2 infection substantially improves after applying phuEGO. **C.** The intersection of phuEGO-derived networks is enriched in known targets for COVID-19.

We hypothesised that if phuEGO is indeed extracting ‘active’ signalling signatures, intersecting the signatures across similar datasets, i.e. those from the same cell type (higgins and stukalov) would result in enrichment of known targets for COVID-19. Indeed, the resulting network, which comprises 85 nodes and 362 edges included 9 known targets, which is 5-fold more than expected by chance (Fisher’s Exact test p-value= 1.60e-04, **Supplementary table S2**). These include SRC kinases LYN, FYN and YES1, of which only YES1 was in the original seed set, and p38 MAPK as well as components of the relevant pathway (e.g. EGFR and BRAF - which is not a known target). Other interesting proteins include RIPK kinases and ROCK1/RhoA which have been previously shown to be advantageous for SARS-CoV2 infection^97^ in relevant genome-wide CRISPR screens.

## Discussion

Signalling processes are very important for the physiological function of cells within their environment and they are highly complex and context-specific. This context-specificity is not captured by the current annotated pathways, which are a result of decades of individual studies and represent the consensus network downstream of individual receptors. It is not practical or feasible to delineate and annotate signalling processes in all possible contexts and conditions in which a cell signalling response occurs; a data-driven approach is therefore needed to identify the active signalling processes from context-specific and unbiased omics data.

Phosphoproteomics data is especially suitable for the study of cell signalling as it measures the signalling state of the cell directly, by providing the signature of phosphorylated proteins and sites in a given moment. As discussed, mapping the data on prior knowledge pathways suffers from literature bias and ignores the context and conditions in which the experiment was done. Conversely, purely data-driven network inference is extremely difficult. This is firstly due to the curse of dimensionality, as no available dataset provides as many data points as phosphosites making the problem unsolvable, and secondly the large understudied signalling space, means that it is anyway very difficult to evaluate methods that do attempt data-driven signalling network inference^34^. Here we present phuEGO, which uses as its basis protein interaction networks^75^, enriched in known signalling regulatory relationships^76–78^. Protein interaction networks are continually becoming more unbiased through systematic efforts such as Bioplex^53^ or HuRI^52^, and therefore they allow us to ground our method on prior knowledge, while at the same time substantially mitigating the severe literature bias that signalling pathways suffer from. As the network is interconnected and no functional units are annotated, our method also allows the data to select the functional modules that are relevant, resulting in functional units that are not restricted by those described by currently annotated pathways, and can better capture cross talk between processes and functional units. In the presented results, we have used a static and universal network for the analysis and the context-specificity of the result stems solely from the data. In addition, while protein interaction networks are far less biased than pathway databases, there still remains certain bias, which is further enhanced by the fact that phuEGO uses semantic similarity to model the edges of the network. This means that poorly annotated nodes would be less preferentially used by the method and those without any annotation are indeed excluded (**Experimental Procedures**). The flexibility of phuEGO, however, means that a user can use any desired network, and this can include for example entirely unbiased predicted networks of functional associations and/or context-specific base networks that take into consideration the transcriptome or proteome of the specific cell line or sample provided, wherever this is available.

The use of network propagation to extract active network modules and signatures is quite well-established^55^ and indeed highly suitable for the study of cell signalling as it simulates signal propagation by the cell through protein interaction networks. However, currently available methods result in large networks that are very hard to interpret. PhuEGO tackles this issue using firstly the semantic similarity to model the edges, to specifically boost the functional signal inherent in the nodes of interest, and secondly applies the local propagation through the ego network deconvolution. The result comprises a much smaller network, organised in distinct functional units/modules that can then be analysed functionally or examined in more detail either independently or viewed at the systems level through supernode links. This can allow the identification of feedback loops, or the prediction of interaction directions, despite the fact that they are not explicitly modelled. Depending on the interest of the user, the parameters can be tuned so that the resulting network is more expanded, albeit noisier, or more specific providing only key signalling processes for the dataset. Providing such precise signalling signatures makes it a lot easier to integrate phosphoproteomics datasets and perform unified analyses as exemplified by the COVID-19 datasets example in this paper. At present, phuEGO performs separate analyses for upregulated and downregulated networks for clarity, but in the future, it is possible to integrate the two to extract single signalling network signatures. In particular, as more functional annotations become available for phosphosites, and the network can include sign and effect of regulatory interactions, phuEGO can provide even more precise signalling signatures from phosphoproteomics data, including increasing the granularity to the phosphosite level, rather than the protein. Finally, the three-layer propagation that phuEGO performs allows us to capture our knowledge with respect to signal transduction and tune the resulting output based on the seeds that we have most confidence in to capture the active signal. In this study we used tyrosine, serine/threonine and non-kinase phosphosites as the three layers, but the method can easily integrate diverse data modalities linking, for example, transcriptomics data, through transcription factor activities, with phosphoproteomics data, through kinase activities and other information.

In conclusion, we present a flexible method, phuEGO, that performs three-layer network propagation on phosphoproteomics data. We show that it is able to boost the signal-to-noise ratio, enrich for functional phosphosites and provide interpretable active signalling network signatures. It is of note that phuEGO performs well both in the high quality, uniformly re-analysed phosphoproteomics datasets in our benchmark^69^ and in the datasets extracted from the original papers. It allows us to better compare and integrate global phosphoproteomics (and other omics) datasets, and potentially other sparse and noisy data types, such as single cell RNAseq. Applying it on five phosphoproteomics datasets derived from cells infected with COVID-19 significantly improved our ability to compare them, and intersecting the two datasets that were collected in A549 cells resulted in significant enrichment of known targets for COVID-19, providing a subnetwork that could point to additional targets. Future improvements of phuEGO include using more unbiased, e.g., predicted, and context-specific networks as its basis and integrating functional annotations of phosphosites to improve the active signalling signature extraction. Overall, phuEGO is a useful tool for the proteomics community and will contribute to the improved study of context-specific cell signalling responses.

## Data availability

All data used in this study is publicly available in the literature, and compiled networks are provided with this work as supplementary tables. phuEGO is freely available as a package through Python Package Index (https://pypi.org/project/phuego/), with source code and documentation hosted on: https://github.com/haoqichen20/phuego.

## Supplemental data

**Supplementary Table 1.** Protein interaction network used as the basis of phuEGO in this study

**Supplementary Table 2.** PhuEGO-derived networks for each of the 5 SARS-CoV-2 datasets used in this study

**Supplementary Table 3.** List of datasets used in the evaluation of phuEGO

**Supplementary Data 1.** Raw output of enrichment analyses performed in each of the 46 datasets used in the evaluation of phuEGO

**Supplementary Data 2.** All networks generated for the SARS CoV 2 analysis from all the datasets included in the study

**Supplementary Figure 1.** Schematic of generation of ego networks **A.** Example of using EGFR as a seed generation to generate the initial ego network **B.** Example of filtering the network using the topological and semantic similarity. **C.** Example of using the Kernel Density Estimation to identify the most topologically and functionally similar nodes to the network.

**Supplementary Figure 2. Schematic of merging ego networks into a supernode network**

**Supplementary Figure 3. Changes in network size at the different steps of phuEGO.**

**Supplementary Figure 4. Characteristics of phuEGO-extracted modules A.** Distribution of number of modules in the datasets tested in this study **B.** Distribution of module sizes generated from the datasets tested in this study.

**Supplementary Figure 5. Evaluation of phuEGO on phosphoproteomics datasets derived from the original publications A.** Comparison of seeds, PCSF, RWR and phuEGO with respect to their ability to rank highly the expected dominant signal, as defined by pathway enrichment analysis. The centre of the circle indicates the relevant pathway ranked first and the perimeter indicates a failure to identify the pathway at any rank. **B.** Association of the overlap coefficient of the phuEGO active signature with the overlap distance of the expected pathways across all pairwise pathway comparisons in our benchmark test. **C.** Phosphosites/nodes retained by phuEGO tend to have a higher functional score indicating an improvement in the signal-to-noise ratio of the active signatures.

**Supplementary Figure 6. Correlation of downregulated phosphosites in SARS-CoV-2 datasets A.** before and **B.** after phuEGO.

