## Supplementary Figures for "phuEGO: A network-based method to reconstruct active signalling pathways from phosphoproteomics datasets"

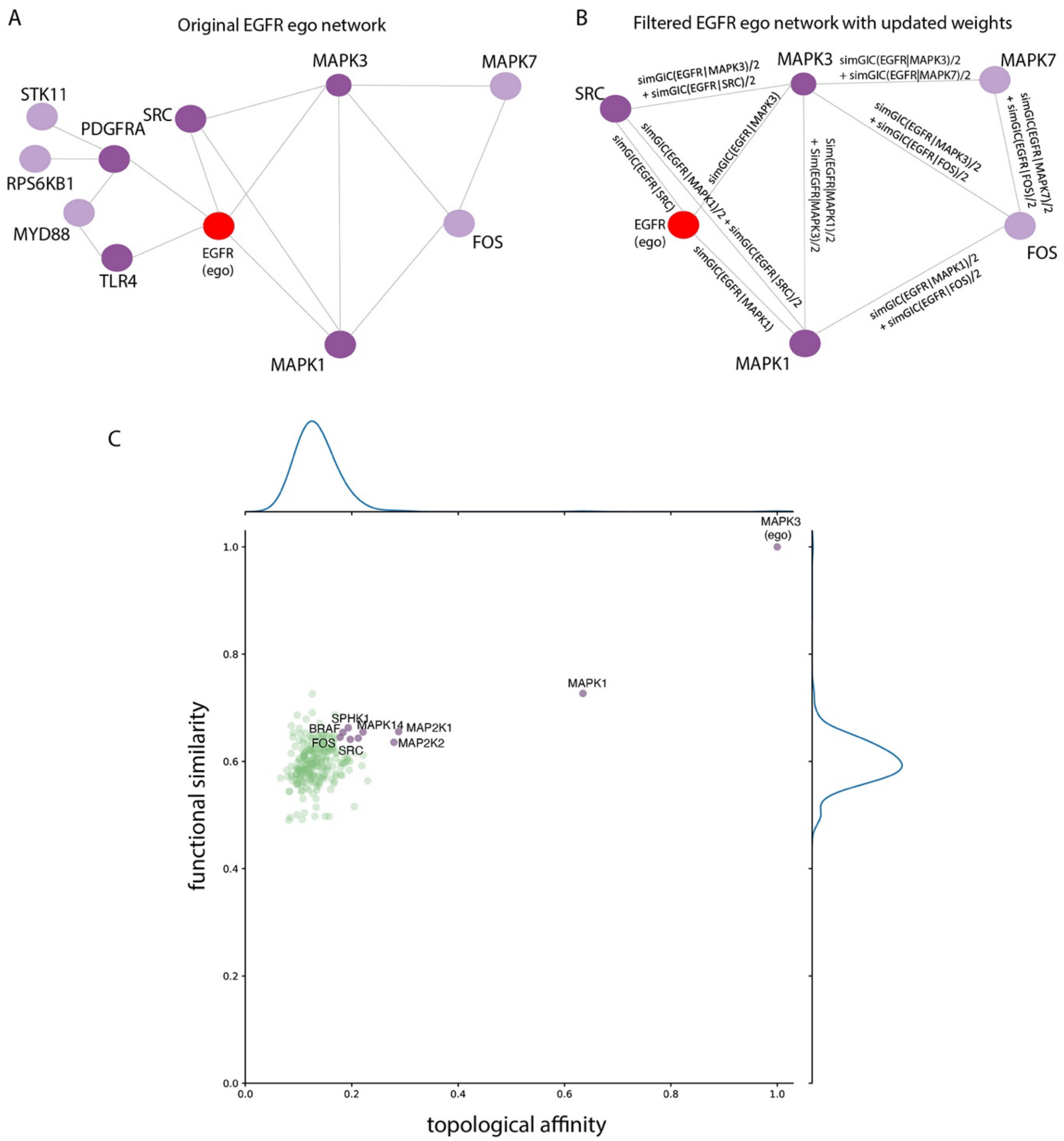


**Supplementary Figure S1. Schematic of generation of EGO networks A.** Example of using EGFR as a seed generation to generate the initial EGO network **B.** Example of filtering the network using the topological and semantic similarity. **C.** Example of using the Kernel Density Estimation to identify the most topologically and functionally similar nodes to the network.

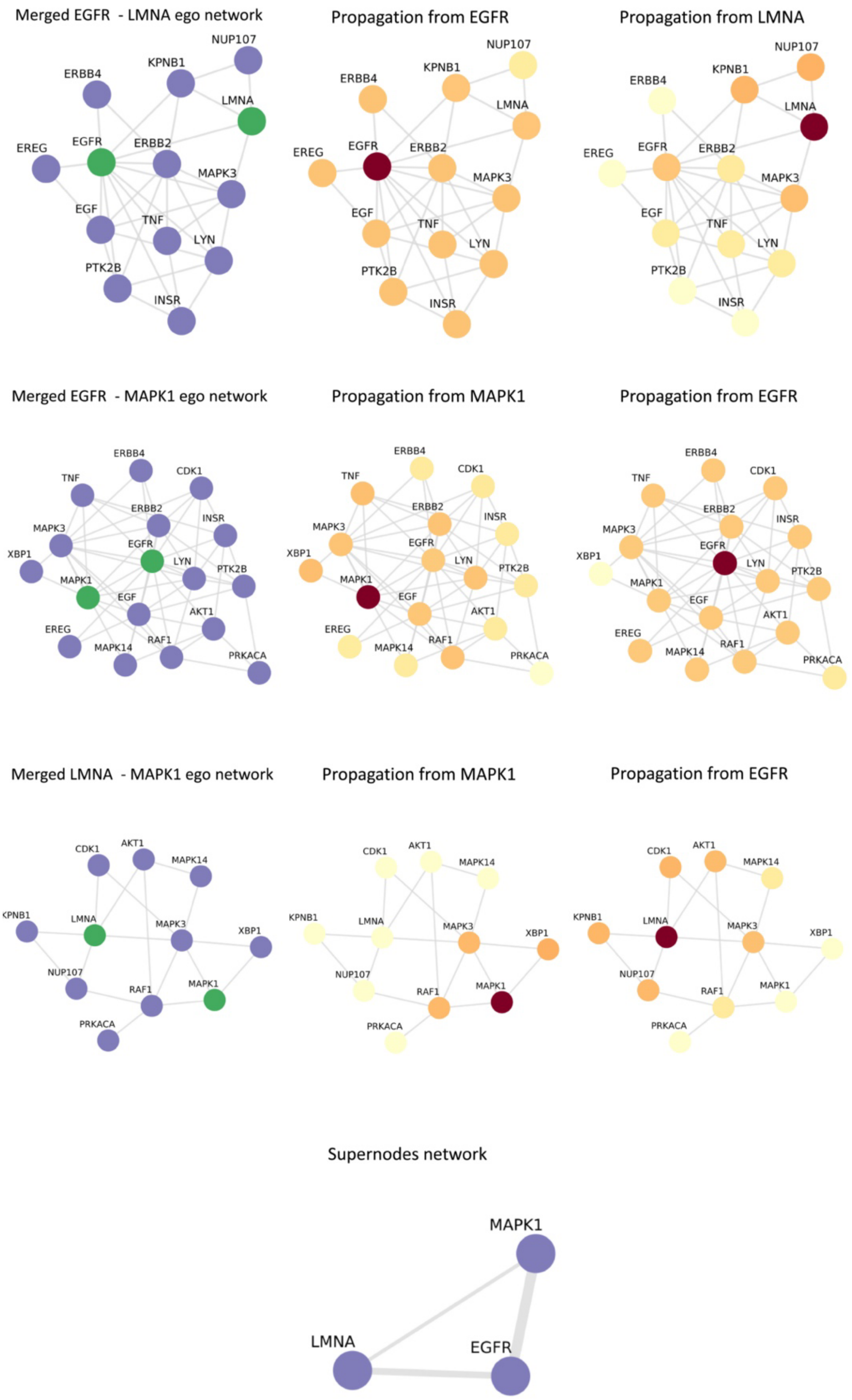


**Supplementary Figure S2. Schematic of merging EGO networks into a supernode network**

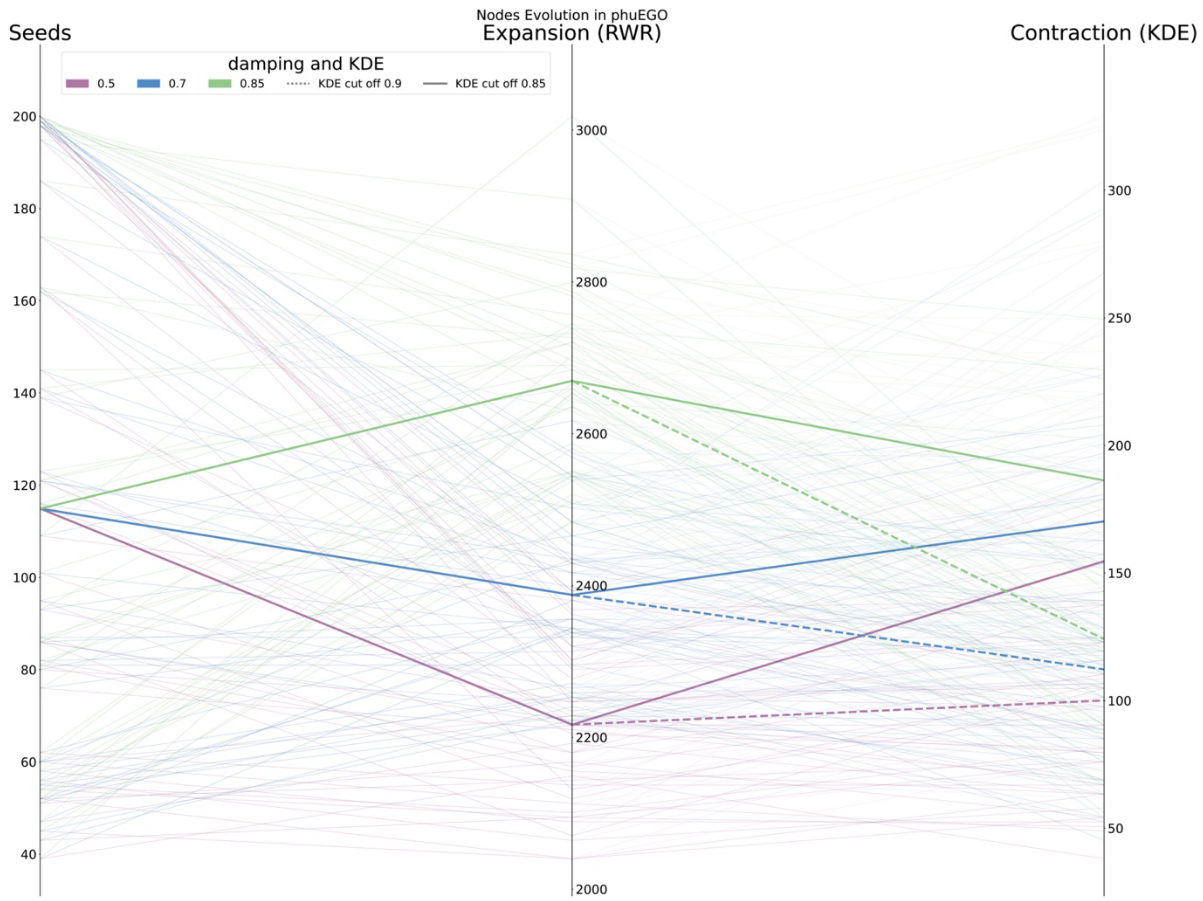


**Supplementary Figure S3. Changes in network size at the different steps of phuEGO.**

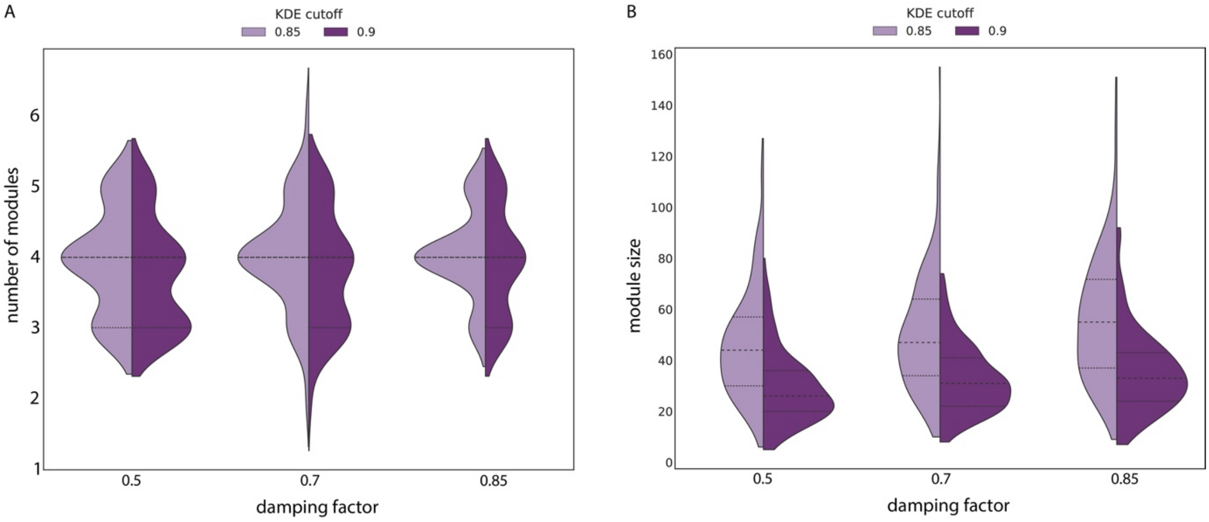


**Supplementary Figure S4. Characteristics of phuEGO-extracted modules A.** Distribution of number of modules in the datasets tested in this study **B.** Distribution of module sizes generated from the datasets tested in this study.

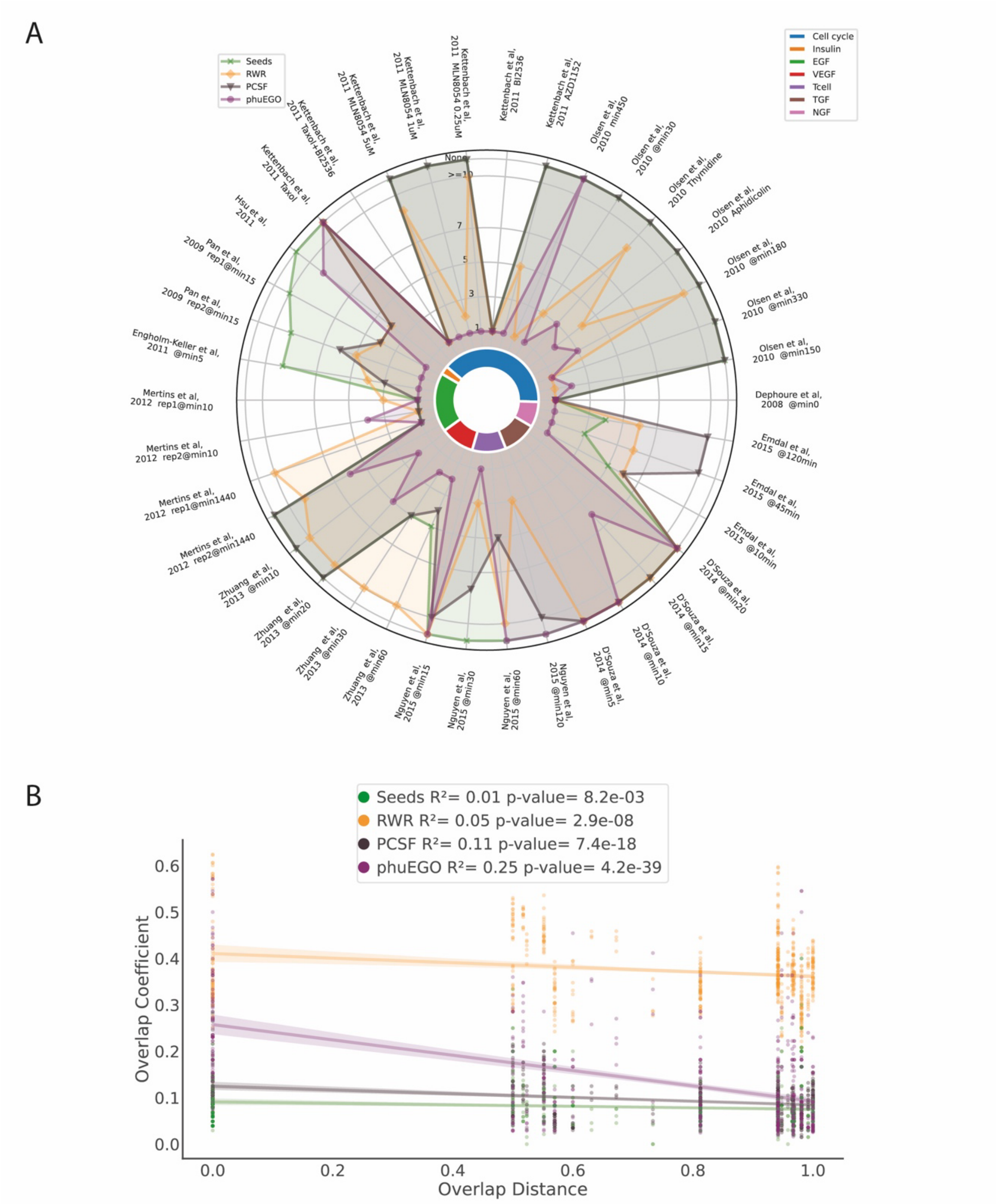


**Supplementary Figure S5. Evaluation of phuEGO on phosphoproteomics datasets derived from the original publications A.** Comparison of seeds, PCSF, RWR and phuEGO with respect to their ability to rank highly the expected dominant signal, as defined by pathway enrichment analysis. The centre of the circle indicates the relevant pathway ranked first and the perimeter indicates a failure to identify the pathway at any rank. **B.** Association of the overlap coefficient of the phuEGO active signature with the overlap distance of the expected pathways across all pairwise pathway comparisons in our benchmark test. **C.** Phosphosites/nodes retained by phuEGO tend to have a higher functional score indicating an improvement in the signal-to-noise ratio of the active signatures.

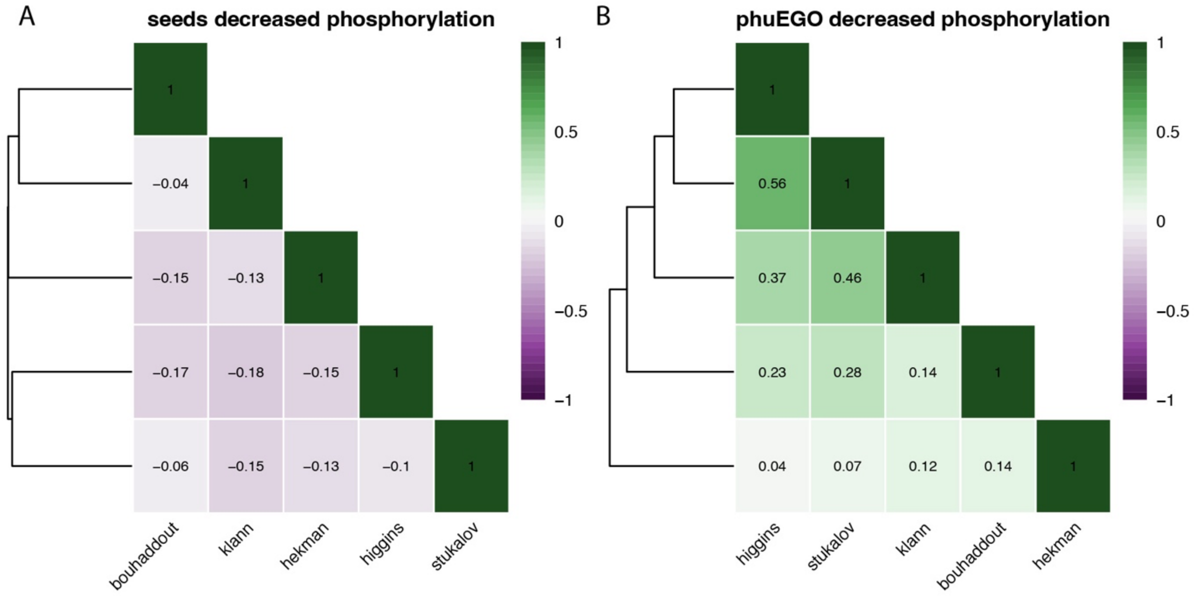


**Supplementary Figure S6. Correlation of downregulated phosphosites in SARS-CoV-2 datasets A.** before and **B.** after phuEGO.
